# Diffusion tubes: a method for the mass culture of ctenophores and other pelagic marine invertebrates

**DOI:** 10.1101/751099

**Authors:** Wyatt L. Patry, Mac Kenzie Bubel, Cypress Hansen, Thomas Knowles

**Affiliations:** Animal Care Division, Monterey Bay Aquarium, Monterey, California, United States of America

## Abstract

The culture of pelagic marine invertebrates, especially the ctenophore *Mnemiopsis leidyi*, has been demonstrated in past studies dating back to the 1960’s, however the *mass* culture of delicate pelagic invertebrates has remained elusive. By using a pair of acrylic tubes and enabling water diffusion between them, we have been able to reliably and cost effectively mass culture several genera of ctenophores (*Pleurobrachia, Hormiphora, Bolinopsis, Mnemiopsis*, and *Leucothea*), one species of siphonophore (*Nanomia*) and one species of larvacean (*Oikopleura*). The simple, compact method is effective enough to support two permanent exhibits of ctenophores at the Monterey Bay Aquarium while minimizing live food culture requirements with the potential to support further investigation of pelagic marine invertebrate ontogeny, ecology and genomics.

## Introduction

Interest in ctenophore culture and other pelagic marine invertebrates as model organisms has surged recently (Howes *et al*., 2014; Jaspers *et al*., 2015, Jaspers *et al*., 2018, Martí-Solans *et al*., 2015; Presnell and Browne, 2019). Basic ontogeny of ctenophores has been observed since at least the turn of the 19th century by Chun, Agassiz and Mayer however maintaining cultures in the laboratory remained elusive until later in the 20^th^ century. Pioneering efforts in developing rearing vessels for pelagic culture were made by Wulf Greve with his invention of the planktonkreisel (Greve 1968 and 1970) and other variations such as the double cuvette design (Greve, 1975). Greve achieved several generations of *Pleurobrachia pileus* in these studies. The kreisel was later formalized by Hamner (1990) for use aboard ships and then adapted for use in public aquariums by the Monterey Bay Aquarium (Raskoff *et al*., 2003).

The method presented here originated at the Monterey Bay Aquarium in July 2015 to support the temporary exhibition, *The Jellies Experience*, and while collaborating with W. E. Browne of University of Miami on *Mnemiopsis leidyi* culture (Presnell *et al*. 2019). After successfully culturing *Mnemiopsis*, we turned our focus to culturing other ctenophores such as *Pleurobrachia bachei* and *Bolinopsis infundibulum*. The young cydippids of these species are so delicate that they do not survive in traditional plankton kreisels. They are susceptible to being damaged by the seawater flow in the tank and sticking to the outflow screen. Additionally, juveniles do not survive in standing seawater dishes due to rapid accumulation of ammonia from waste, and high mortality from being transferred into new seawater. Therefore, we needed a rearing tank with minimal or no mechanical driven water flow, no outward pressure on the outflow screen, and the ability to passively exchange new seawater. We experimented with rearing tanks featuring seawater flow just outside of the outflow screen, such that seawater could passively exchange without creating outward pressure on the screen. While these methods were effective at eliminating buildup of nitrogenous wastes and minimizing forces on juvenile ctenophores caused by seawater flow, the tank shapes were not appropriate for the swimming and feeding behavior of the animals. In traditional rectangular aquaria, juveniles were observed actively consuming prey at or near the surface and then sinking to the bottom, where significant biofouling occurs. Initially we designed and constructed a cylindrical tank with more vertical space for feeding, minimal biofouling surface area with mesh covering the bottom and secured in a pseudo-kreisel such that water flowed up through the bottom. This initial design proved somewhat effective, however juveniles still spent significant time contacting the bottom mesh. An improved design providing even more vertical space, while reducing the volume of the tank was implemented. Our final version utilizes a double cylinder configuration in which an elongated inner cylinder is surrounded by an outer cylinder ‘sleeve’, affectionately referred to as a “cteno-tube” in the lab.

Over the past four years, we have experimented with this new double cylinder rearing tank design and have refined a protocol for raising fragile pelagic ctenophores. We observed a number of physical parameters including; feeding rates, biofouling, and analyzed fluid flow rates within the ‘cteno-tube’ in relation to adult ctenophore yield. Here we present these results, additional variations on the double-cylinder method and provide a best practices protocol for the mass culture of pelagic ctenophores and other gelata.

## Method

### Materials and design

Diffusion tubes were constructed using two 0.9 m long acrylic tubes. The outer and inner tubes had volumes of ~35 L and 28 L with inner diameters of 21.59 cm and 20 cm respectively, so that one could be placed inside the other (an inner and outer cylinder); each cylinder has a thickness of 0.3175 cm (Fig. 1). The bottom of the inner cylinder was fitted with 55 μm nylon screening (Model# M55 PentairAES, Inc.) using silicone sealant (Dowsil 795 or 999-A) (Fig. 1c). This inner cylinder was then placed on a 3 cm tall riser made of rigid plastic mesh (Model# N1020 PentairAES, Inc.) such that the top of the inner cylinder rises above the outer cylinder (Fig. 1a). The outer cylinder was glued to a ~1 cm thick square acrylic baseplate, 15-20 cm^2^, using acrylic cement (Weld-On #16 Fast Set solvent cement). Rigid PVC tubing, 0.635 cm inner diameter, (Part# 48855K41, McMaster-Carr) was placed in the outer cylinder, between tubes, providing water flow near the bottom. Filtered seawater (5 μm) was pumped through the tubing at rates of 1.1, 2.2 and 4.5 Liters per minute (Lpm) based on ½, 1, and 2 theoretical turnovers of total cylinder volume per hour. Three identical pairs of diffusion tubes were constructed and placed on a wet table with recirculating filtered seawater (5 μm) such that water overflowed out of the outer cylinder and on to the table (Fig. 2a). Diffusion between tubes through the 55 μm nylon screening was observed by the addition of dye (McCormick blue food color) to the incoming water. In the 4.5 Lpm treatment, flow was such that a slight suction vortex pulled the ctenophore eggs into the bottom screen which prevented hatching. To resolve this, a PVC pipe tee fitting (Part# 4881K47, McMaster-Carr) was added to the bottom of the inflow tube to reduce water velocity and divert the incoming water away from directly underneath the screen, which proved successful (Fig. 2b).

**Figure 1.**
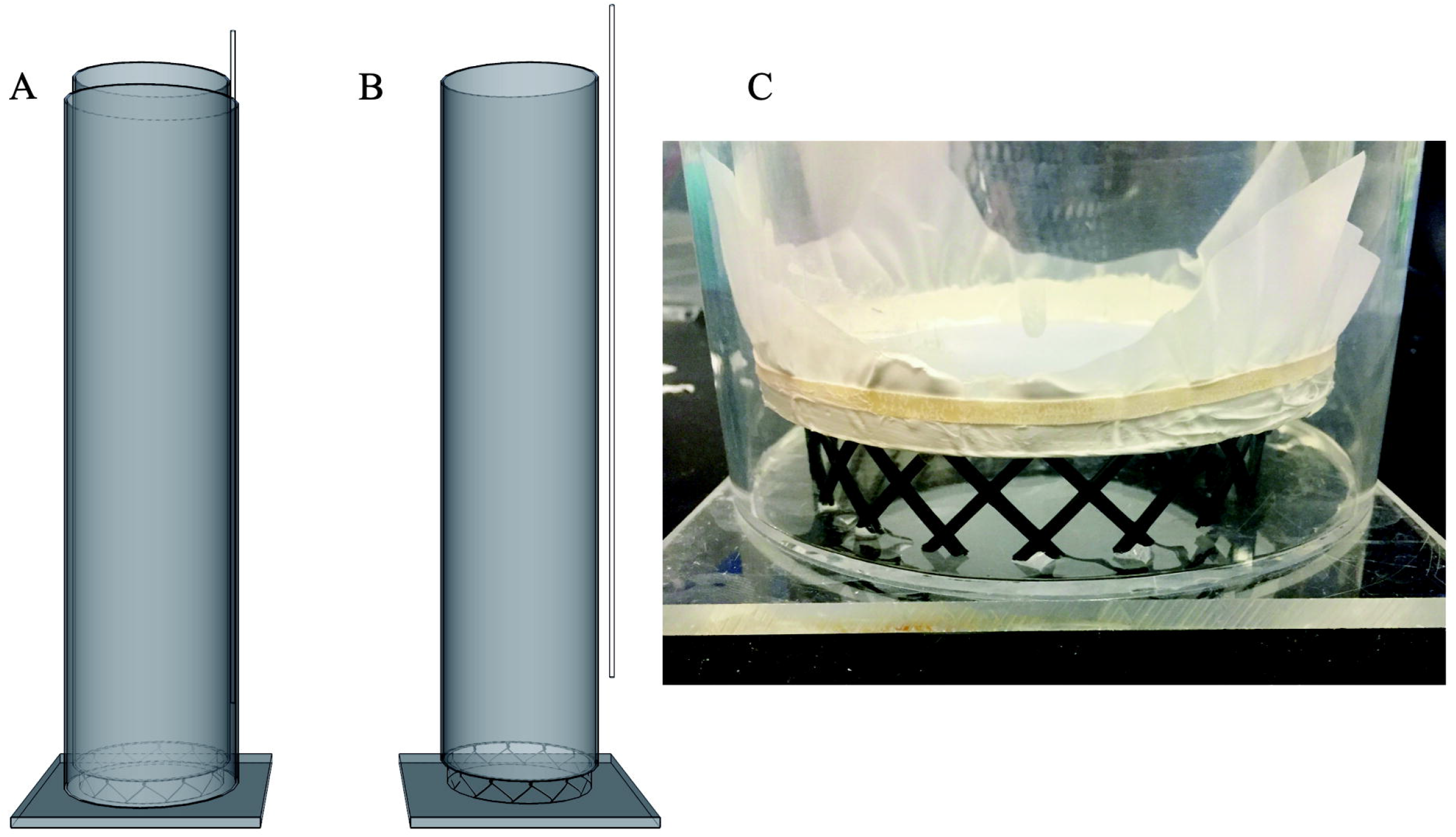
Rendering and construction of the diffusion tubes. (A) both tubes are visible, with supply tubing at the top right. (B) outer tube has been removed to reveal the inner tube resting on the riser, made with rigid plastic mesh. Water supply tubing is entirely visible on the right, note how tubing does not reach to the bottom (−15 cm above the bottom plate) to avoid entrainment of eggs into the screen. (C) bottom portion of tube shown with plastic riser. A rubber band may be used to secure the mesh for gluing with silicone sealant, excess mesh can be cut off. Care should be taken when lowering the inner tube to avoid tearing the mesh on the riser. Riser should be cut to the exact diameter of the inner tube so the mesh does not rest on the riser.

**Figure 2.**
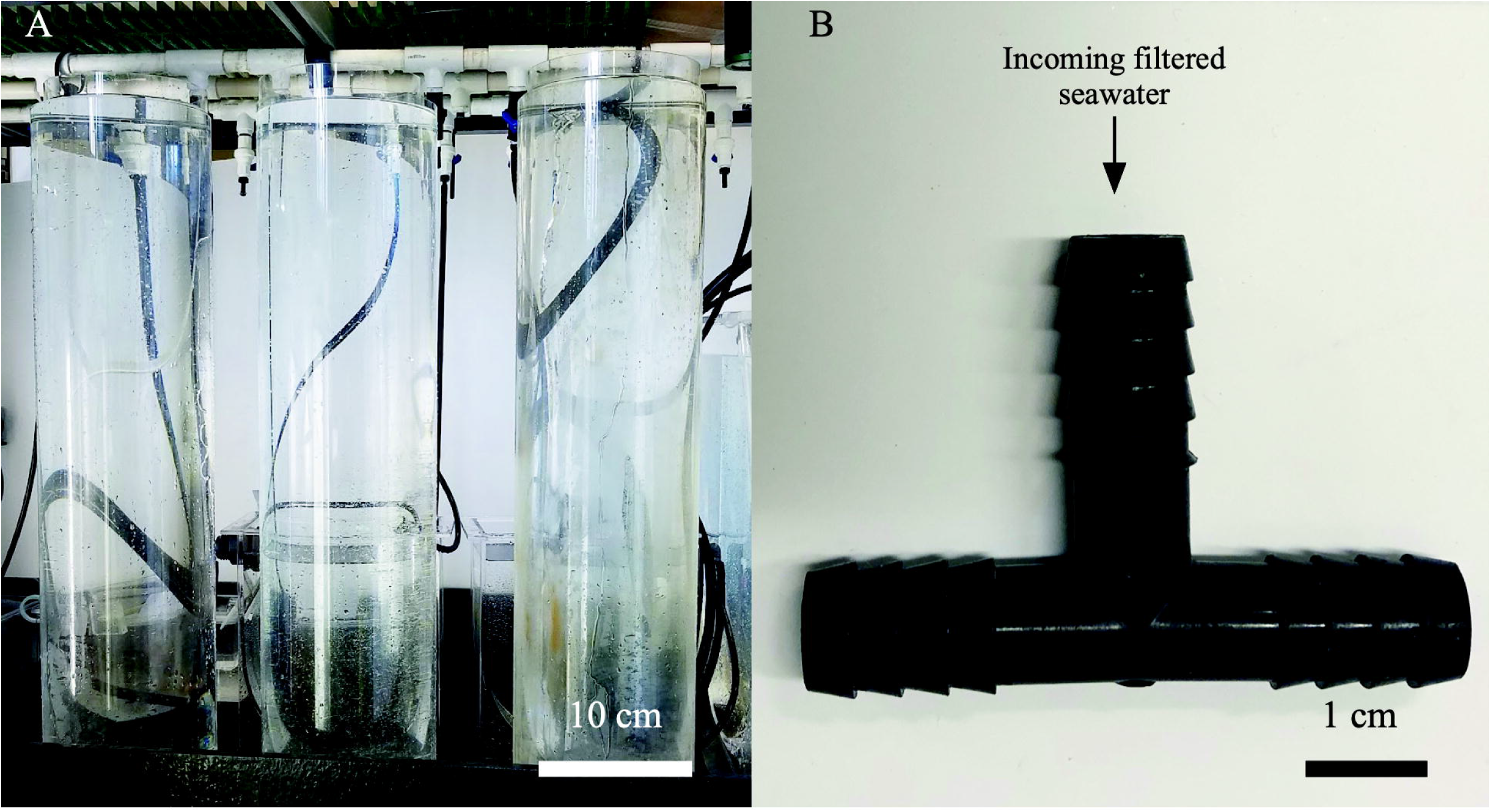
Experimental setup. (A) three sets of diffusion cylinders on a wet table. (B) Tee fitting used to prevent entrainment of eggs on screen.

### Characterizing physical parameters

A set of probes (Hach HQ40d with LDO probe) for measuring pH, dissolved oxygen, and temperature was deployed within the inner cylinder of each set of tubes and reads were taken every 12.7 cm at seven discrete depths. Tubes were allowed to equilibrate for at least 24 hours prior to testing and the probes were lowered slowly to read depths using a simple pulley system. A minimum settling time of two minutes at each depth was necessary to take accurate reads of each parameter. We also used a dye to observe mixing/stratification within the tube and found that the inflow tube caused flow disturbance resulting in water primarily diffusing in from the opposite side of the tube inflow. In order to account for discrepancies caused by the inflow tube’s position on one side vs. the other, probe readings were established twice, once closest to the inflow tube and once on the opposite side.

### Methods of spawning and culturing

Three adult *Hormiphora californensis* were placed in each inner cylinder (Fig. 3a). To induce spawning (Fig. 3b), the tubes were shrouded in a black, opaque cover for total darkness overnight, followed by 2 hours of bright light the next morning (Kessil A360 Tuna Blue LED aquarium light placed ~5 cm above the inner cylinder) (Pang & Martindale 2008). When eggs were observed throughout the water column, usually ~12 hours post spawn or 24 hours after setup, the adults were removed from the tubes using a spoon attached to a short pole or by slowly swirling them to the surface.

**Figure 3.**
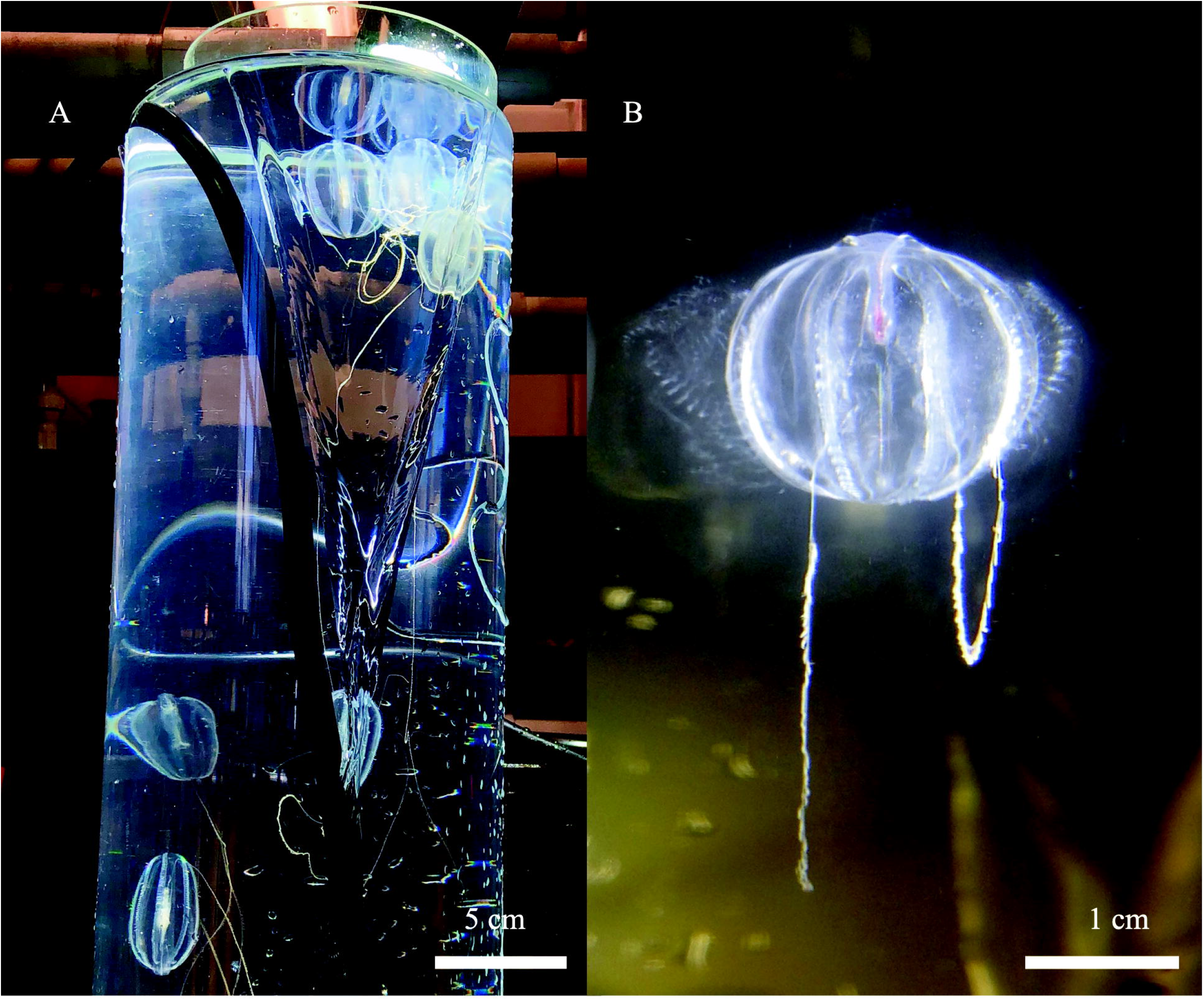
Spawning ctenophores in the tubes. (A) *Hormiphora californensis* spawning in the inner cylinder with a bright LED light. (B) *Pleurobrachia bachei* releasing sperm from the meridional canals beneath the ctene rows.

Hatching of the eggs was observed 12 - 48 hours post spawning. Juvenile cydippids were fed on the first day post hatch (dph). Each tube was given a standardized concentration (~30 nauplii mL^-1^) of live *Parvocalanus crassirostris* (Reed Mariculture Inc.) nauplii mixed with live algae *(Isochrysis galbana, Rhodomonas lens, Dunaliella tertiolecta*, or *Tetraselmis chuii*, Florida Aqua Farms, Inc.) in a 1:1 ratio, both cultured in the lab. *P. crassirostris* nauplii were added at the surface of the tube water column for six weeks using the following regimen: 25 mL 3 times per week (Sunday, Tuesday, Thursday) for the first 14 days, increasing to 50 mL for the next 14 days, and 100 mL for the last 14 days.

The ctenophores were observed throughout the water column and sampled once a week for six weeks to be photographed and measured. For photo documentation and measurement, 15 - 30 juvenile cydippids were randomly pipetted out of the water column down to 30 cm depth and transferred to a glass crystallizing dish filled with filtered seawater for photographing under a microscope (Zeiss Stereo Zoom V16, Canon EOS Rebel T5i). Each photograph was scaled and measured using ImageJ 1.52A (Schneider et al., 2012).

When cydippids reached ~0.5 - 1 cm in diameter (~6 weeks post hatch) they were transferred using widened disposable polyethylene pipettes (tips cut off, Transfer pipets 13-711-7M, FisherBrand) or small polypropylene beakers (50 - 100 mL Nalgene Griffin low form beaker) from the diffusion tubes to kreisel tanks (Raskoff et al., 2003) modified to provide for better longevity of ctenophores. The kreisel screens were replaced with a polyethylene mesh with 4.0 mm openings (Model #N1670 PentairAES, Inc.). Slot openings in the supply boxes were reduced in width using a single piece of 4 mm corrugated plastic (high-density polyethylene and polypropylene) instead of two 6 mm pieces (Model #24244SC CorrugatedPlastics.Net). Pumps were set to provide ~ 10 liters per minute (Lpm) of volumetric flow to the lower supply box and ~7.5 Lpm to the upper box (Model DCW-2000 Jebao, Inc.) using flow meters (Blue-White Industries, Ltd, Model #F-44500L-8). The upper box provided ~0.8 Lpm of filtered seawater (5 μm) at 13°C. The diet was diversified to include 1 - 2 mysid shrimp *(Mysidopsis bahia*, Aquatic Indicators, Inc.) per comb jelly, usually ~50 per kreisel and 200 mL of *P. crassirostris* adult copepods, fed daily around 1200 PST.

### Statistical analysis

Mean length data was analyzed using R(R Core Team, 2017). Growth rates were adapted to the data using the R package Growthcurver(https://cran.r-project.org/package=growthcurver). Descriptive statistics, Kruskal-Wallis ANOVA and Wilcoxon pairwise comparisons were made using packages dplyr(https://CRAN.R-project.org/package=dplyr). ggpubr(https://rpkgs.datanovia.com/ggpubr/) and ggplot2(http://ggplot2.tidyverse.org/).

## Results

### Physical parameters and depth profiling

#### Temperature

Under the highest turnover rate (4.5 Lpm), the water column exhibited a significant thermocline within 12-24 cm of the surface. With a surface temperature between 16-17°C, the surface layer was approximately 3-4 degrees warmer than the water below 24 cm in depth (Fig. 4a). This was in agreement with dye observations, which indicated slower turnover in the top half of the water column under the 4.5 Lpm treatment. Although the other two treatments exhibited less dramatic thermoclines, the mean surface temperature exceeded the mean bottom temperature by at least 1.5 °C in all three treatments. Temperatures reported are the mean of the measurements taken on the two opposite sides of the inner cylinder, nearest and farthest from the inflow line. No significant difference was found between the near and far side measurements.

**Figure 4.**
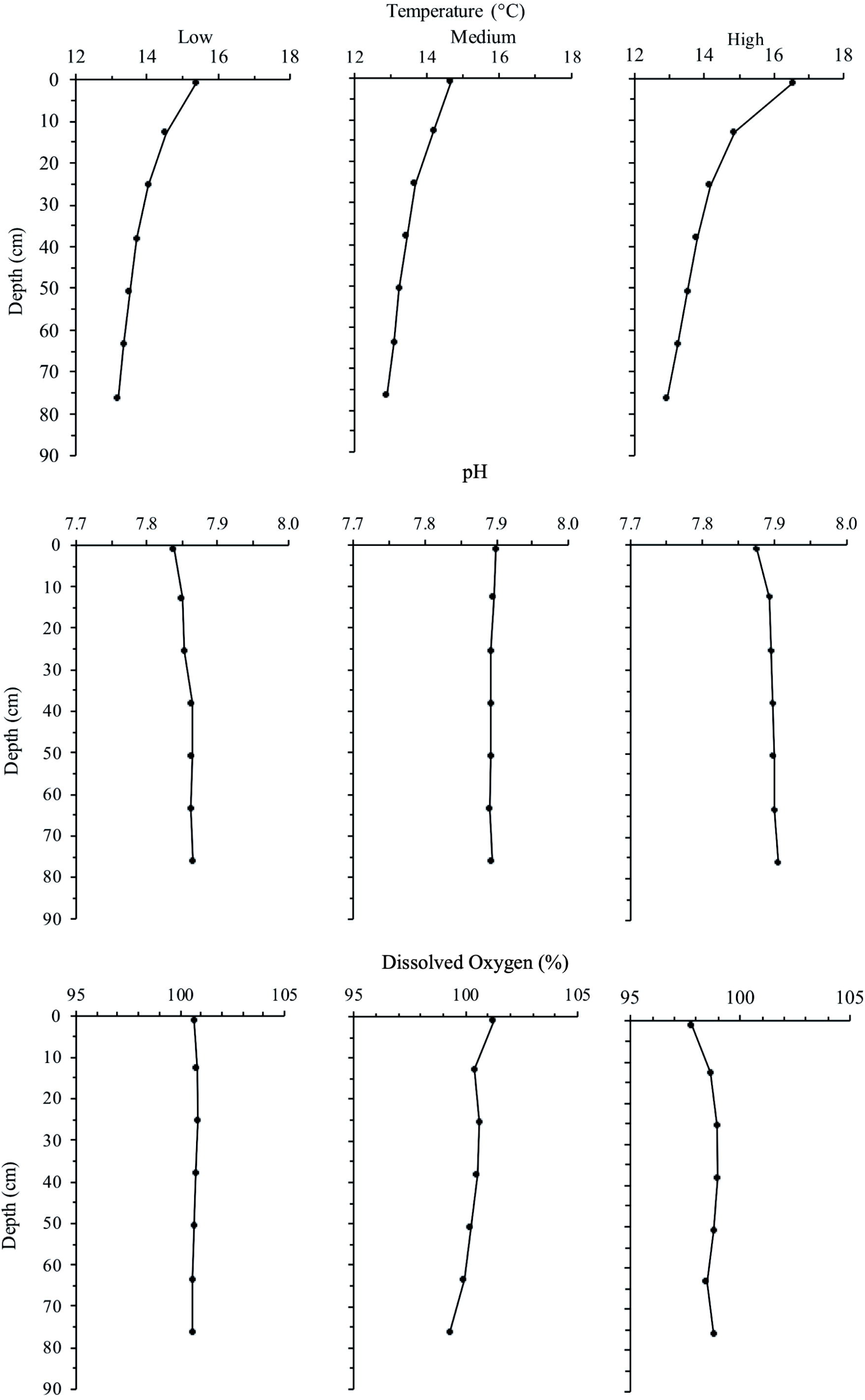
Depth profiles of each treatment for temperature, pH and dissolved oxygen. Line is the mean of the two measurements taken on opposing sides of the inner tube.

#### pH

pH was the most consistent of all parameters measured, ranging between 7.84 and 7.90 across all flow rates and depths, with the low flow treatment yielding the lowest pH at the surface of the water column (Fig. 4b). The low and high flow treatments exhibited slightly decreased pH at the surface layer.

#### Dissolved Oxygen

Dissolved oxygen measurements exhibited no significant differences with regard to depth, flow rate, or temperature. The high flow treatment showed slightly decreased dissolved oxygen at the surface layer in conjunction with decreased pH, however the measurements were within the standard error of the probe (Fig. 4c).

#### Culturing success

All three treatments were successful in raising >50 *H. californensis* over a six-week period. The high flow (4.5 Lpm) treatment yielded the most cydippids (89), followed by medium flow (1.1 Lpm) (87) and low flow (2.2 Lpm) (75). High flow and low flow treatments reported the following growth curves respectively, K = 33.705, N = 0.026, r = 0.574 and K = 33.759, N = 0.013, r = 0.635. Growthcurver failed to find a best fit for the medium flow treatment. The mean cydippid length of each treatment varied little for the first 4 weeks of growth (each p > 0.07) and then greatly increased during the fifth week (Fig. 5). Cydippids in the high flow treatment had the highest mean lengths at the end of the experiment (Wilcoxon signed-rank test, p = 0.00054 and 1.3 E-6, low vs. high and medium vs. high respectively), while the medium treatment had the smallest mean lengths (Fig. 6).

**Figure 5.**
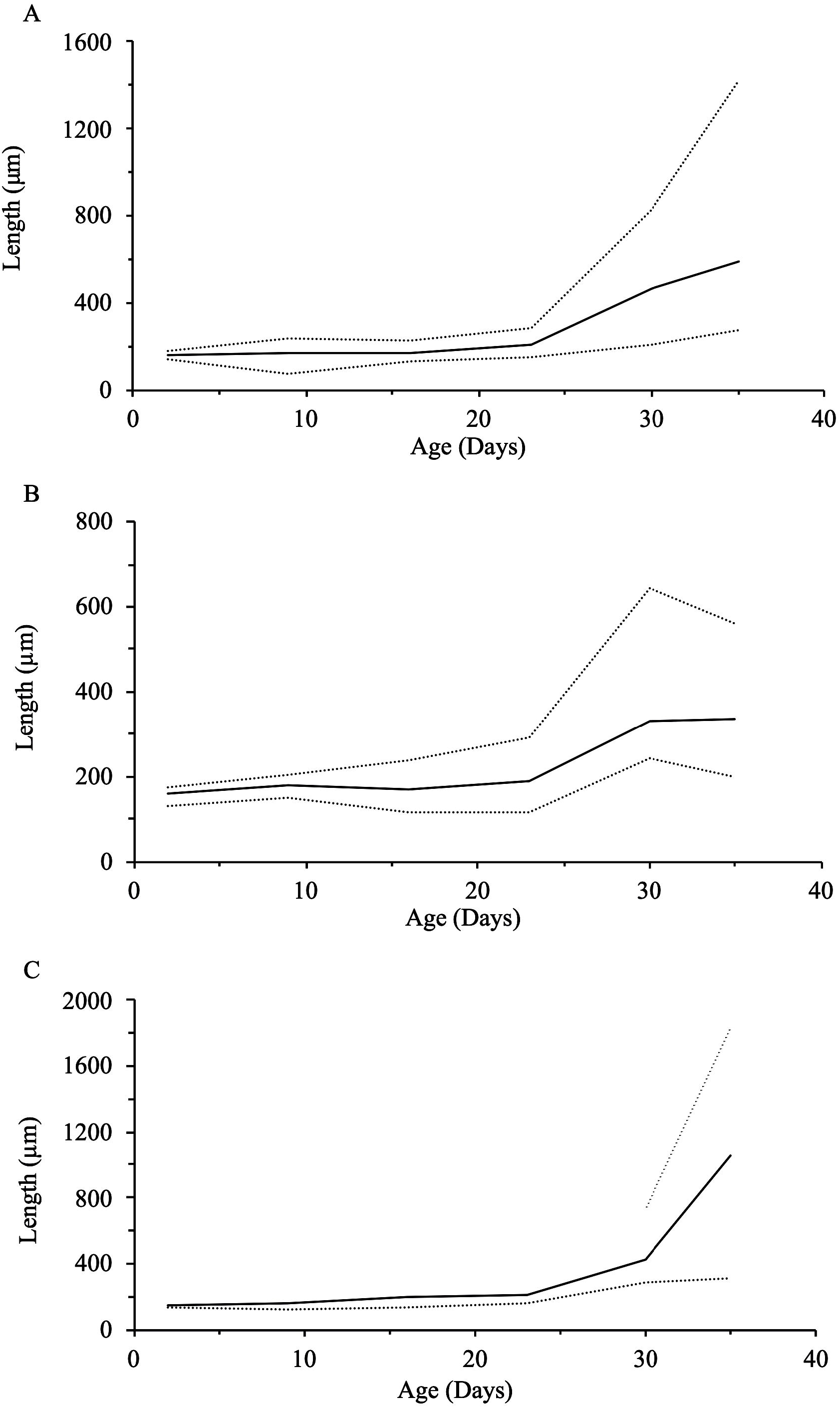
Total length measured over 35 days. Average (solid line), minimum, and maximum (dotted lines) total lengths of *H. califomensis* in low (A), medium (B), and high (C) flow treatments.

**Figure 6.**
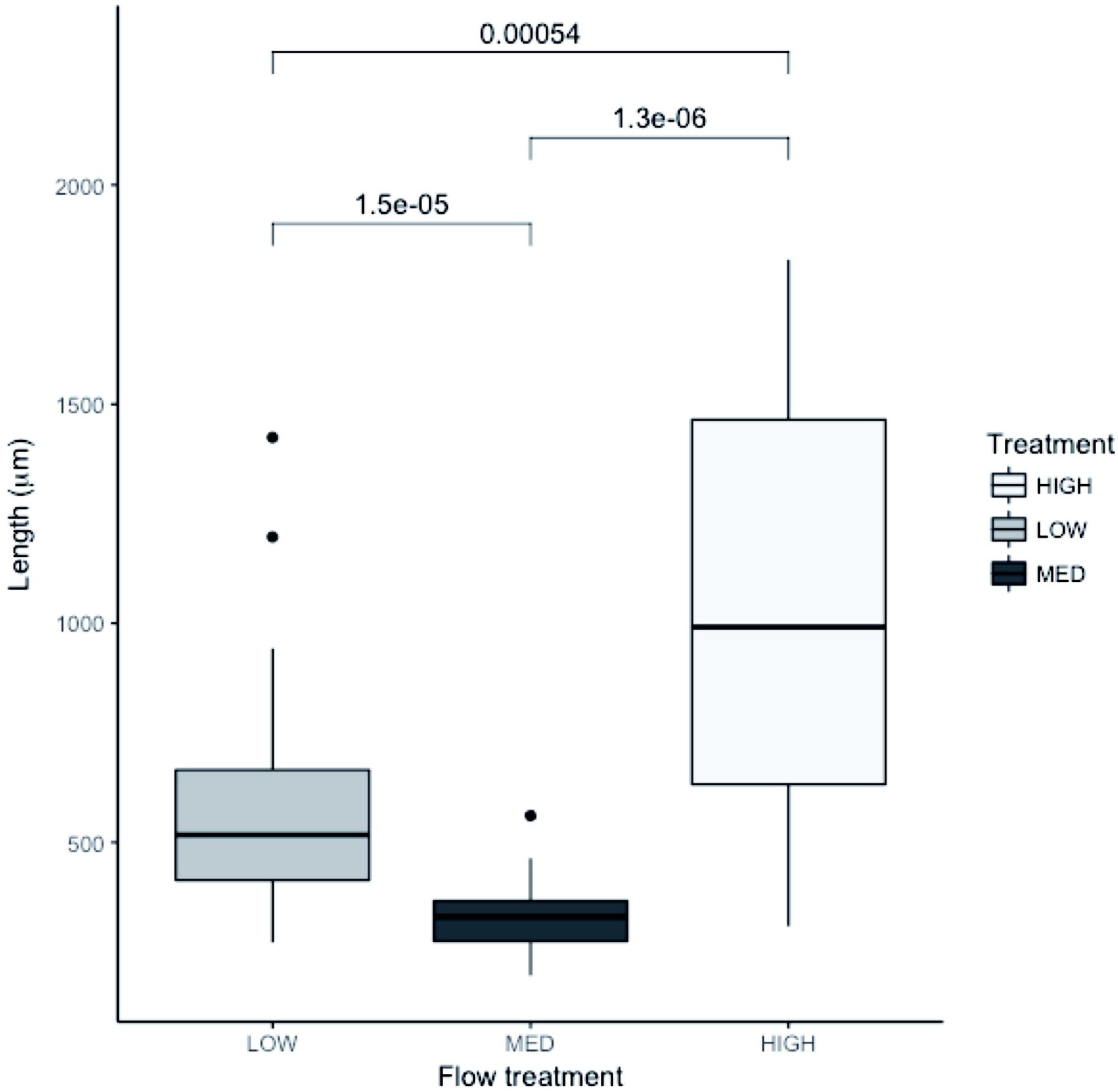
Pairwise comparison of means for low, medium, and high flow treatments at week 6. The high and medium flow treatments ended with significantly greater mean lengths than the low flow treatment. The high flow treatment had the greatest mean lengths.

## Discussion

Our method utilizes a cylindrical tank shape which maximizes the ratio of water volume to the surface area of tank bottom, which has several important effects. By minimizing the bottom biofouling area, contact with these fouled surfaces is greatly decreased, vertical swimming or migration space for juveniles is maximized and a thermocline is produced where the juvenile cydippids can feed furthest from the tank bottom. Another key feature of our design is the passive exchange of seawater through the mesh which generates much lower water velocities than in a traditional plankton kreisel. These lower flow rates allow delicate juvenile ctenophores to develop in a very gentle environment while still facilitating the exchange of clean filtered seawater. In contrast to traditional aquaria for gelatinous zooplankton, our method provides a superior pelagic environment for young ctenophores and other gelata larvae that is inexpensive and easily replicable.

Using a higher turnover rate generated a more pronounced vertical thermocline and resulted in higher survival and significantly faster growth rates among the treatments. It is not clear why the medium flow outperformed the low flow treatment; perhaps further replication would provide resolution. Greater concentrations of *P. crassirostris* were observed at the surface of the inner cylinder in the high flow treatment and this appeared to be optimal for ctenophore feeding, helping to keep cydippids actively feeding near the surface and away from settled particulates at the bottom of the inner cylinder. Fouling was limited to minor algal compaction on the bottom mesh and cydippids did not appear to spend significant time there or become impacted.

Overall, fewer *H. californensis* were produced in this experiment than previous and subsequent trials with the diffusion tubes. We believe this is due to the restricted quantities of food used. When feeding quantity was increased from 50 mL to 100 mL at week 5 there was a notable increase in growth rate across treatments. It is also possible that lower yields may reflect using fewer parents (n = 3) than previous/subsequent trials (n = 10). Ultimately, we adopted a feeding regime that doubled the amounts of copepod nauplii (see Conclusions). An F4 population was achieved shortly after the experiment using 10 parents of the F3 population, yielding >600 cydippids.

Further development of this method has the potential for increased efficiency and enable use of the tube method with new species of pelagic gelata. Changing cylinder height and width may allow for more variation in thermocline profiles and vertical space, however taller tubes will require more volumetric flow to the outer cylinder. For larger ctenophore species diffusion tubes with larger diameters may have increased utility, for example *Leucothea pulchra* was successfully cultured to F2 using diffusion tubes 2 m high and 0.35 m diameter with a flow of ~4 Lpm (Fig. 7). *Bolinopsis infundibulum* and *Nanomia bijuga* were grown using an inner cylinder measuring 0.9 m high and 0.25 m diameter (outer cylinder was 0.9 m high and 0.30 m diameter) (Fig. 8). Changing the mesh size may impact diffusion rates between the two tubes, thermocline formation and perhaps be better fitted to a particular target species. For example, a 22 μm mesh was originally used for *P. bachei* but we found the mesh fouled quickly. When replaced with 55 μm mesh, diffusion and flow performance were enhanced and fouling no longer a problem.

**Figure 7.**
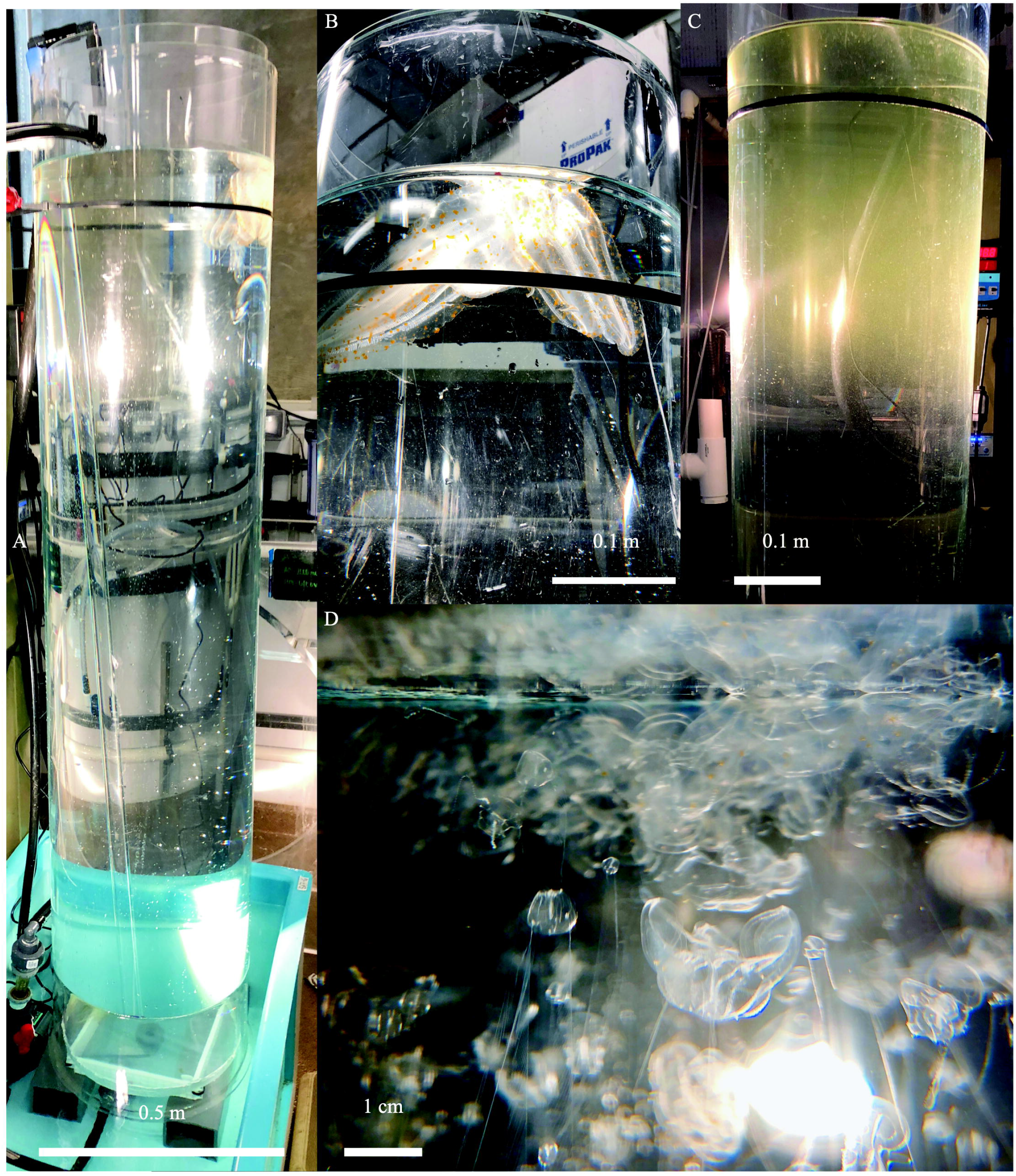
Taller diffusion tube setup. (A) Two meter tall diffusion tubes for rearing *Leucothea pulchra* sitting inside of a 200 L reservoir. (B) three large adults were spawned to produce the FI generation. (C) thermocline visible at the top of tube, 9 dph (D) various srages of FI larval *L. pulchra* visible at the surface of the inner tube, 21 dph.

**Figure 8.**
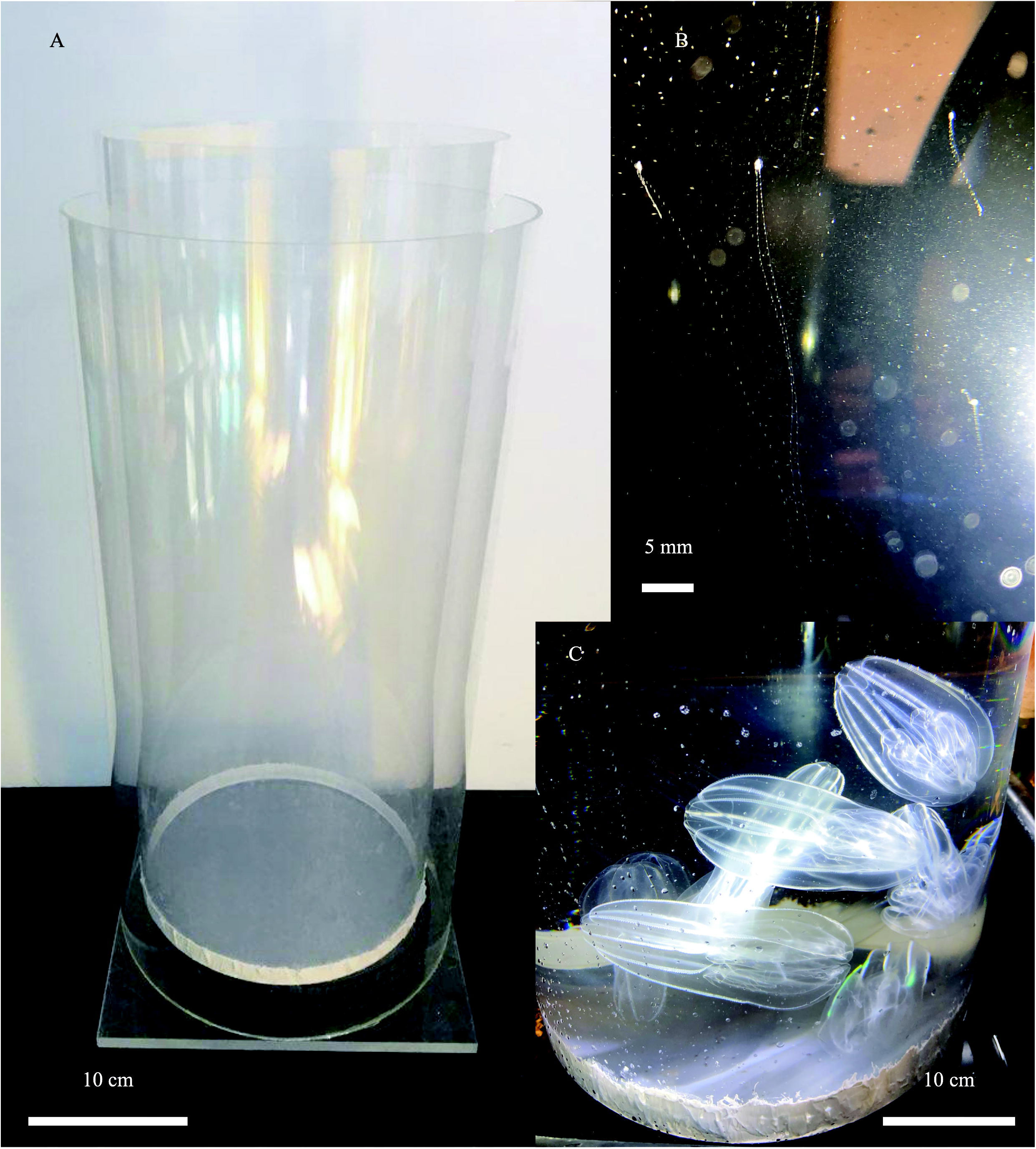
Shorter and wider diffusion tubes setup. (A) 30 cm wide diffusion tubes used for *Bolinopsis infundibulum* and *Nanomia bijuga* culture. (B) larval *N. bijuga* at 35 days post settlment visible near the top of the tube (C) mature *B. infundibulum* spawning under bright light, the wider and shorter tube makes it easier to place and remove the adult lobates.

## Conclusions

After four years of trials culturing various gelata using different sizes of the diffusion tubes, we derived a standard procedure for mass culture of ctenophores from 1-10 parental adults using 1 m diffusion tubes (*Hormiphora* and *Pleurobrachia*) and a feeding regime based on results yielding >100 adult ctenophores in 4-5 weeks was adopted as follows:

I. To start a culture from wild collected adults, use 1 - 10 adult ctenophores depending on desired genetic diversity. Adults should be fed a fish or mysid meal 24 hours prior to spawning and again 1 hour before the dark period begins (Presnell et al., 2019).
II. To induce spawning, place all adults in an equilibrated diffusion cylinder (for inner diameter 20 cm, a flow rate of 4.5 Lpm), shrouded in opaque material to ensure complete darkness for a 4-hour period (or overnight). Uncover the tube and expose to bright lighting for at least 2 hours (Presnell et al., 2019).
III. When adults have spawned and eggs are visible in the water column, gently remove them from the tube using ladles or dishes.
IV. Eggs hatch within 12-48 hours, first feeding should ideally take place within the first 24 hours of hatching to ensure that cydippids are able to begin feeding immediately.
V. Feeding schedule: see Table 1.
VI. At 4-5 weeks post hatching the cydippids may be moved to a pseudo-kreisel or kreisel tank. If cydippids have not achieved a total body size of 0.5 - 1 cm diameter by this time, their residence in the tube may be increased to as long as 6 weeks. However uniformly slow growth rates are a sign of a weak hatch, not enough food and/or significant disruption in water flow/diffusion through the tube. As residence time increases in the tube, so will impaction of the screen.

Gelatinous zooplankton require physically stable conditions to grow *en masse*. Our tube design maximizes water exchange through an efficient and gentle diffusion profile, reduced fouling, increased residence time of live prey, and significant total water turnover with very little or no mechanical turbulence. The diffusion cylinder method can be a critical tool for laboratory use in the cultivation of a variety of pelagic gelata species and thus contribute to a range of research in these fragile organisms on population genetics, zooplankton plastivory, behavior and ecology.

## Supporting information

Feeding Table 1

## Acknowledgements

We thank Dr. W.E. Browne for his enthusiastic collaboration, comments on draft manuscript and support of ctenophore culturing here at the aquarium. The DEEP-C project for creating a joint ctenophore project that introduced us to Dr. Browne and other ctenophore researchers. Shannon Johnson for her comments on the draft manuscript. John Negrey for instrument calibration and support. John Hoech, Paul Clarkson, Marcus Zevalkink, Sarah Halbrend, Evan Firl, the Drifters gallery staff and volunteers that contributed their time and resources to this project.

## Funding

Funding was provided by the Monterey Bay Aquarium Foundation.

## Notes

#### Summary of Updates

Figures formatted, Figure 6 revised. Table 1 created.

