## Supplementary material for "Diffusion tubes: a method for the mass culture of ctenophores and other pelagic marine invertebrates": Feeding Table 1

Table 1: Feeding schedule:

| Time | Week 1 | Week 2 | Week 3 | Week 4 | Week 5 |
| --- | --- | --- | --- | --- | --- |
| Frequency | 3 times per week | 3 times per week | 3 times per week | 3 times per week | 3 times per week |
| Copepod nauplii | 50 mL | 100 mL | 200 mL | 400 mL | 400 mL |
| Adult copepods | None | 50 mL | 100 mL | 200 mL | 400 mL |

Feeding Notes:

a) Feeds may be skipped when copepods remain in the tube uneaten.
b) When copepods remain uneaten in the tube, live algae (*Isochrysis*, *Tetraselmis*, *Rhodomonas* etc.) may be added to the next feed as an equal volume to the feed. This helps keep the remaining feed enriched and growing.
c) Feed may be placed in a beaker and slowly dripped into the top of the inner tube using a drip emitter or similar device (DYNALON clamp, part #670715)
